# Efficient approximations for stationary single-channel Ca^2+^ nanodomains across length scales

**DOI:** 10.1101/2020.01.16.909036

**Authors:** Y Chen, C Muratov, V Matveev

## Abstract

We consider the stationary solution for the Ca^2+^ concentration near a point Ca^2+^ source describing a single-channel Ca^2+^ nanodomain, in the presence of a single mobile Ca^2+^ buffer with one-to-one Ca^2+^ binding. We present computationally efficient approximants that estimate stationary single-channel Ca^2+^ nanodomains with great accuracy in broad regions of parameter space. The presented approximants have a functional form that combines rational and exponential functions, which is similar to that of the well-known Excess Buffer Approximation and the linear approximation, but with parameters estimated using two novel (to our knowledge) methods. One of the methods involves interpolation between the short-range Taylor series of the buffer concentration and its long-range asymptotic series in inverse powers of distance from the channel. Although this method has already been used to find Padé (rational-function) approximants to single-channel Ca^2+^ and buffer concentration, extending this method to interpolants combining exponential and rational functions improves accuracy in a significant fraction of the relevant parameter space. A second method is based on the variational approach, and involves a global minimization of an appropriate functional with respect to parameters of the chosen approximations. Extensive parameter sensitivity analysis is presented, comparing these two methods with previously developed approximants. Apart from increased accuracy, the strength of these approximants is that they can be extended to more realistic buffers with multiple binding sites characterized by cooperative Ca^2+^ binding, such as calmodulin and calretinin.

**STATEMENT OF SIGNIFICANCE:** Mathematical and computational modeling plays an important role in the study of local Ca^2+^ signals underlying vesicle exocysosis, muscle contraction and other fundamental physiological processes. Closed-form approximations describing steady-state distribution of Ca^2+^ in the vicinity of an open Ca^2+^ channel have proved particularly useful for the qualitative modeling of local Ca^2+^ signals. We present simple and efficient approximants for the Ca^2+^ concentration in the presence of a mobile Ca^2+^ buffer, which achieve great accuracy over a wide range of model parameters. Such approximations provide an efficient method for estimating Ca^2+^ and buffer concentrations without resorting to numerical simulations, and allow to study the qualitative dependence of nanodomain Ca^2+^ distribution on the buffer’s Ca^2+^ binding properties and its diffusivity.

## I. INTRODUCTION

Some of the most fundamental physiological cell processes such as synaptic neurotransmitter release, endocrine hormone release, muscle contraction and cytotoxic immune cell response are directly and quickly triggered by the Ca^2+^ influx into the cytoplasm (1-4). Due to the diversity of Ca^2+^-controlled cellular processes, intracellular Ca^2+^ signals are localized in time and space to allow selective activation of specific reactions (2-5). This localization is maintained in part by intracellular Ca^2+^ buffers, which absorb most of the Ca^2+^ influx soon upon its entry into the cell (6,7). In the context of secretory vesicle exocytosis, local Ca^2+^ concentration elevations around individual Ca^2+^ channels or clusters of channels are termed Ca^2+^ nano- or micro-domains (4,8). Although Ca^2+^ concentration can be measured experimentally using Ca^2+^ sensitive dyes, inherent physical limitations pose challenges for optical Ca^2+^ imaging on small temporal and spatial scales relevant for vesicle exocytosis and other processes controlled by local Ca^2+^ elevations. Therefore, mathematical and computational modeling has played an important role in the study of vesicle exocytosis and other cell processes activated by localized Ca^2+^ signals (8-15). In particular, these computational studies were instrumental in showing that local Ca^2+^ elevations form and collapse very rapidly in response to channel gating. This suggests that quasi-stationary solutions of the reaction-diffusion equations describing Ca^2+^ influx, diffusion and binding to intracellular Ca^2+^ buffers may achieve sufficient accuracy in estimating Ca^2+^ concentration in the vicinity of a Ca^2+^ channel, obviating computationally expensive solutions of partial differential equations describing buffered Ca^2+^ diffusion (16,17). Several of such stationary approximations have been introduced in the early works of Neher, Stern, Keizer, Smith and others (14,18-26), most notably the Excess Buffer approximation (EBA), the Rapid Buffering approximation (RBA), and the linear approximation (LIN) (see Table 1). These approximations proved quite useful in understanding the properties of Ca^2+^ nanodomains and their dependence on the properties of cell Ca^2+^ buffers, and widely used in modeling studies (9,14,21,27-30). However, most of the previously developed approximations have two limitations: (1) their accuracy is restricted to specific regions in buffering parameter space, and (2) they have been developed for simple, one-to-one Ca^2+^-buffer binding, and are hard to extend to more realistic buffers that have multiple Ca^2+^ binding sites (31).

**Table 1.** Previously established single-channel equilibrium Ca^2+^ nanodomain approximations. For each method, only the free buffer concentration expression is shown, since the non-dimensional Ca^2+^ concentration can be found from the Ca^2+^ conservation condition (Eq. 11), with the exception of terms in 2^nd^ order EBA, which are shown in (20). Note that LIN and EBA become identical in the limit ν>>1. RBA approximations valid up to orders O(1) and O(*λ*) are denoted as 1^st^-order RBA (or simply RBA) and 2^nd^-order RBA (RBA2), respectively. Two lowest orders of the Padé method are denoted Padé for the 1^st^ order case, and Padé2 for the 2^nd^ order case. For Padé2, the expressions for parameter-dependent rational function constants *A*_l,2_ and *B*_l,2_ are given by the solution of a 4^th^ order polynomial equation, which are closed-form but too lengthy to show explicitly here (32).

| Method | Free buffer concentration, $b(r)$ | Conditions | References |
| --- | --- | --- | --- |
| LIN | $1 + \frac{q}{r} \left[ \exp(-r/\sqrt{q\lambda}) - 1 \right]$ | Linearization around $c_\infty = 0$ | (14,18,20,24,25,37) |
| EBA | $1 + \frac{1}{\nu r} \left[ \exp(-r\sqrt{\nu/\lambda}) - 1 \right] + O(1/\nu^2)$ | $\lambda \gg 1, \nu \gg 1, \frac{\lambda}{\nu} = O(1)$ | (14,20,38) |
| IBA | $\eta \left[ \frac{r}{1+\eta r} + \frac{\nu r^2}{(1+\eta r)^3} + \frac{2\lambda}{(1+\eta r)^4} \right] + O(\lambda^2)$ | $\lambda \ll 1, \nu \ll 1, \frac{\lambda}{\nu} = O(1)$ | (20) |
| RBA | $1 - \frac{1}{2q\nu} \left[ 1 + \frac{q}{r} - \sqrt{\left(1 + \frac{q}{r}\right)^2 - \frac{4\nu q^2}{r}} \right] + O(\lambda)$ | $\lambda \ll 1, \nu = O(1)$ | (14,19-21,23,27) |
| RBA2 | $b_{RBA}(r) + 2\lambda\eta \left[ \left(1 + \frac{r}{q}\right)^2 - 4\nu r \right]^{-2} + O(\lambda^2)$ | $\lambda \ll 1, \nu = O(1)$ | (20) |
| Padé | $1 - \frac{q}{r + \left[ \sqrt{q(q+8\lambda)} + q \right] / 2}$ | Series interpolation | (32) |
| Padé2 | $\frac{r^2 + A_1(\lambda, \nu, \eta)r + A_2(\lambda, \nu, \eta)}{r^2 + B_1(\lambda, \nu, \eta)r + B_2(\lambda, \nu, \eta)}$ | Series interpolation | (32) |

Here we present several improved approaches allowing to better approximate single-channel Ca^2+^ nanodomains with more accuracy and for a wider range of model parameters. One of these approximation methods is based on matching the coefficients of short-range Taylor series and long-range asymptotic series of the nanodomain Ca^2+^ distance dependence using a simple *ansatz*. Although this method has already been used to obtain Padé (rational function) nanodomain approximations (32), we show that significant improvement can be achieved in some parameter regimes using alternative interpolants that are similar in their functional form to EBA and LIN approximants. Similar *ansatz* can also be extended to buffers with multiple binding sites (work in progress). Apart from the local-series interpolation approach, we also present a different class of methods based on global optimization of a relevant functional with respect to parameters of the same *ansatz* that we use with the series interpolation method, which have superior accuracy in certain parameter regimes, as demonstrated below.

## II. METHODS

### II.1 Single-channel Ca^2+^ nanodomain equation

Following prior work, we will consider a Ca^2+^ buffer whose molecules possess a single active site that binds a Ca^2+^ ion according to the reaction

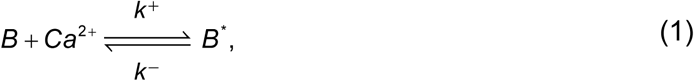

where *B* and *B*^*^ are the free buffer and Ca^2+^-bound buffer, respectively, and *k*^+^/*k*^−^ are the Ca^2+^-buffer binding/unbinding rates. We consider a semi-infinite diffusion domain bounded by a flat plane containing point Ca^2+^ channel sources. Following previous modelling studies (19,20,26), we will assume Dirichlet boundary conditions on the outer boundary representing the background concentrations for Ca^2+^ and buffer in the bulk of the cell cytoplasm, and zero flux boundary condition on the flat boundary representing the cell membrane. Although this neglects Ca^2+^ pumps and exchangers along the flat boundary, numerical simulations show that qualitative agreement with more accurate models is retained under this assumption. The reflection symmetry along the flat boundary allows to extend the domain to the whole space, while doubling the source strength. Assuming mass-action kinetics, this yields the following reaction-diffusion system in ℝ^3^ (19,20):

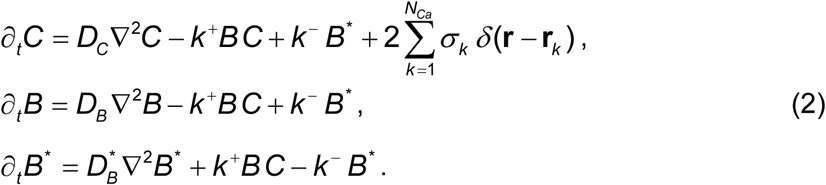

Here *C, B* and *B*^*^ represents concentrations of Ca^2+^, free buffer and Ca^2+^-bound buffer, respectively, with diffusivities *D*_*C*_, *D*_*B*_, and 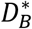. In the source term, *N*_*Ca*_ denotes the number of Ca^2+^ channels, and the source strengths are given by *σ*_*k*_ = *I*_*Ca,k*_/(*z F*), where *I*_Ca,k_ are the amplitudes of individual open Ca^2+^ channels located at positions **r**_*k*_, *F* is the Faraday constant, and *z*=2 is the valence of the Ca^2+^ ion. We note that the point-like channel assumption introduces inaccuracy at small spatial scales commensurate with the channel pore width of several nanometers. The impact of finite channel diameter and volumetric Ca^2+^ clearance was considered in a different type of single-channel stationary solution derived for the endoplasmic reticulum Ca^2+^ channel in (16). The two linear combinations of Eq. 2 that cancel the reaction terms yield the conservation laws for the total Ca^2+^ and total buffer concentrations:

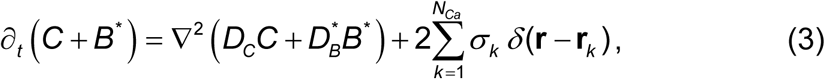

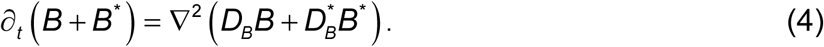

We now consider the steady state of this system, where the conservation laws for Ca^2+^ and buffer reduce to (20,21,23,26,33,34):

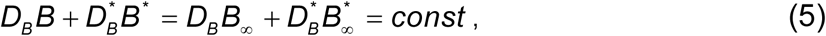

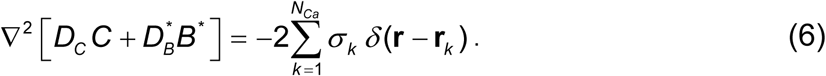

Our approach is somewhat more general than prior modeling work in that we do not assume that buffer mobility is unaffected by Ca^2+^ binding. Given our simplifying assumptions on the domain geometry and boundary conditions, Eq. 6 has an exact solution:

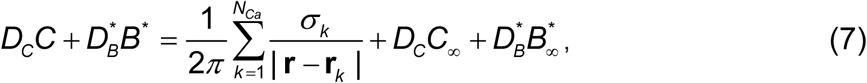

where *C*_∞_ and *B*_∞_ are the background Ca^2+^ and buffer concentrations infinitely far from the channel, which are in equilibrium with each other:

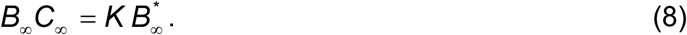

Here *K* = *k*^−^/ *k*^+^ is the buffer affinity, equal to the Ca^2+^ concentration at which half the buffer is bound at steady state. Conservation laws allow to eliminate two variables, and we choose to retain the equilibrium unbound buffer concentration as the remaining unknown:

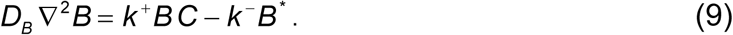

We will now non-dimensionalize these equations similar to the method of Smith et al (20) (see also (32)), rescaling Ca^2+^ by the buffer affinity: *c*=*C*/*K, c*_∞_=*C*_∞_/*K*. However, we normalize the buffer concentration by its background value *B*_∞_ instead of total concentration. This will simplify analytic results, with many expression formally unchanged whether or not *c*_∞_=0 (see Table 1). Note also that in this case a very simple relationship holds between background concentrations of Ca^2+^ and bound buffer: Eq. 8 yields 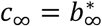. We will consider the case of a single channel at the origin, and re-scale the spatial coordinate (**r** /*L* → **r**) using the scale parameter that depends on the strength of the Ca^2+^ current, which simplifies the source term in Eq. 7 (20):

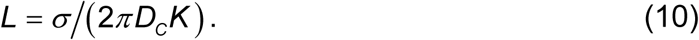

Recalling that 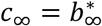, we obtain the following non-dimensional form of free buffer dynamics given by Eq. 9, and the conservation laws, Eqs. 5, 7:

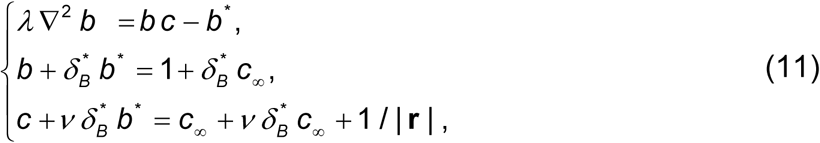

where the four non-dimensional model parameter are (with *L* given by Eq. 10):

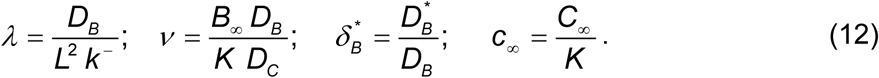

Here *λ* is the dimensionless buffer diffusion coefficient (denoted as *ε*_*b*_ in (20)), which quantifies the diffusion rate relative to the rate of Ca^2+^ binding and influx, while *ν* (denoted as 1/*μ* in (20)) represents the overall buffering strength at rest, given by the product of the resting buffering capacity (*B*_∞_/*K*) and the relative buffer mobility (*D*_B_/*D*_C_). In this non-dimensionalization, unbuffered Ca^2+^ solution corresponds to *ν*=0 and has a particularly simple form, *c* = 1/|***r***| + *c*_∞_. For the sake of simplicity, we will also use the following auxiliary parameters:

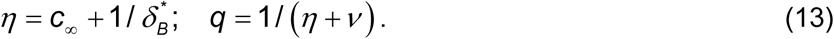

This allows to specify the problem using only 3 parameters, either {*λ, ν, η* } or {*λ, q, η* }. Eliminating bound buffer and Ca^2+^ concentrations using the two conservation laws in Eqs. 11, the free buffer equation takes on a simple form:

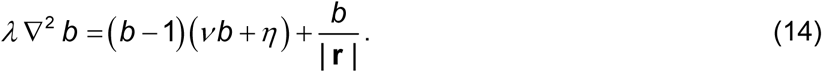

Ca^2+^ concentration can be obtained from the solution of Eq. 14 using the Ca^2+^ conservation law in Eq. 11, which can be simplified to the following intuitive form:

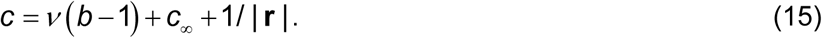

For *b*<1, Ca^2+^ concentration is reduced in proportion to the buffering strength parameter *ν*, as expected. The conservation laws in Eq. 11 along with the physical constraints *c*≥0, *b*^*^≥0, *c*_∞_≥0 imply *a priori* bounds

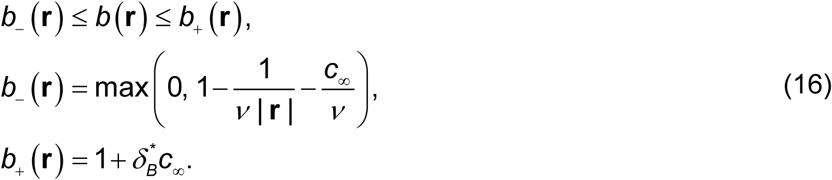

Solutions satisfy the following boundary conditions (here and below, we denote *r* =| **r** |):

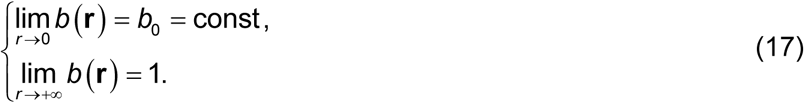

where the value of buffer at the source location, *b*_0_, is unknown *a priori*. As is rigorously proved in Appendix 3, Eq. 14 has a unique solution in a suitable function space, and this solution is spherically symmetric. Therefore, Eq. 14 may be reduced to

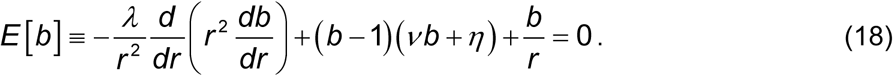

Although Eq. 18 superficially resembles the Lane–Emden-Fowler equations (35), it has no local Lie symmetries allowing analytical solution. Further, it is not of Painlevé type (36), despite its simple algebraic form. We carried out the numerical solution of Eq. (18) using the relaxation method and the shooting method, cross-validating the results of these two methods. For certain extreme values of model parameters, accurate numerical solution is computationally intensive.

We note that the chosen non-dimensionalization is identical to the one in (20,32) in the case of binding-invariant buffer mobility 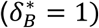 and zero background Ca^2+^ concentration (*c*_∞_=0). In the case *c*_∞_≠0 there is a simple equivalence with the non-dimensionalization in (20,32); indicating variables and parameters in the latter work with the hat symbol, this equivalence reads:

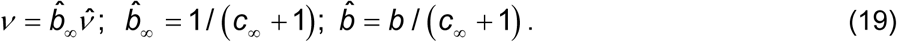

Since in biologically relevant cases *c*_∞_≈0, in numerical results shown further below we focus on the special case *c*_∞_=0, *η*=1. However, all results of this study were validated using a range of *η* values, ensuring that our conclusions are not sensitive to the value of *η.*

One of the contributions of early modeling efforts was the development of accurate analytical approximations of the solution of Eq. 18. They allow avoiding computationally expensive integration of reaction-diffusion equations while retaining considerable accuracy (20,32). These approximations are summarized in Table 1. As was previously shown in (20), the Excess Buffer approximation (EBA), Rapid Buffering approximation (RBA) and the Nearly Immobile Buffer approximation (IBA) represent asymptotic expansions in either *λ, ν*, or *μ*=1/*ν*, while the linear approximation (LIN) is obtained by an *ad hoc* linearization around the free unbuffered point-source solution, *b* = 1, *c* = 1/*r* + *c*_∞_. In contrast, the Padé approximation is based on a series matching method explained in detail below (32). We note that only 2^nd^ order RBA and Padé approximations are comparable in accuracy to the approximants presented in this work. Since [Ca^2+^] is uniquely determined by the buffer concentration through the conservation laws (Eq. 11), [Ca^2+^] estimation accuracy is only shown in the final summary and comparison of all approximations (see Figs. 5 and 6). We note that accurate estimation of free buffer concentration can be as important as the knowledge of the corresponding Ca^2+^ concentration, since it helps in the understanding of cell Ca^2+^ homeostasis, and in interpreting the results of Ca^2+^ imaging, which requires quantifying Ca^2+^ binding to exogenously applied fluorescent Ca^2+^ buffers (2,3,5,8).

**Figure 1.**
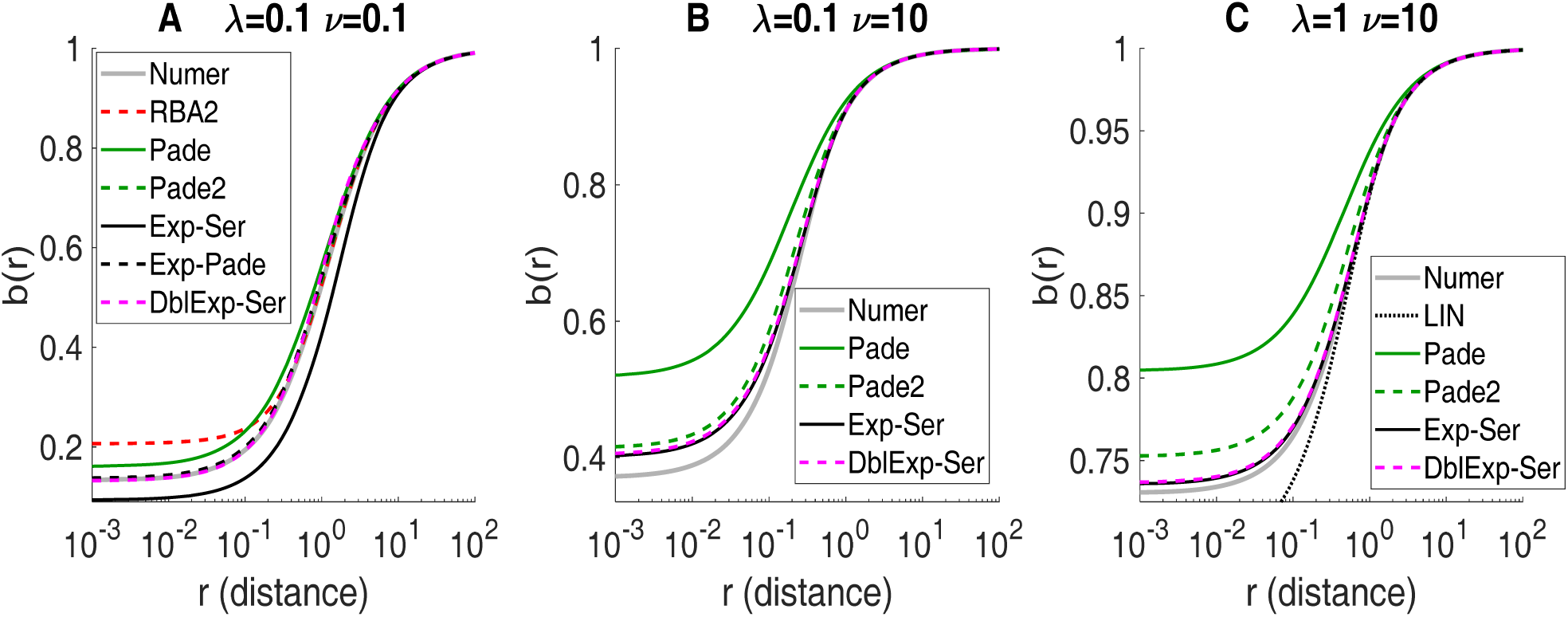
Equilibrium nanodomain buffer concentration approximations obtained using the series interpolation method: 1^st^-order Padé (*green*), 2^nd^ order Padé (Padé2, *dashed green*), Exp-Ser (*black*), Exp-Padé, (*dashed black*), and DblExp-Ser (*dashed magenta*). Also shown for comparison is RBA2 (A, *dashed red*) and Linear approximation (C, *dotted black*). All panels show free dimensionless buffer concentration as a function of distance from the Ca^2+^ channel, for 3 distinct choices of model parameters *λ* and *ν*, as indicated in the panel title, with *η* =1. *Grey curves* show the accurate numerical solution. In (A), DblExp-Ser, Padé2 and Exp-Padé are indistinguishable from the numerical solution on this scale. Note that Exp-Padé does not yield a solution for *ν* >*η* =1 (B,C). In (A), DblExp-Ser curve shows the real part of Eq. 26.

**Figure 2.**
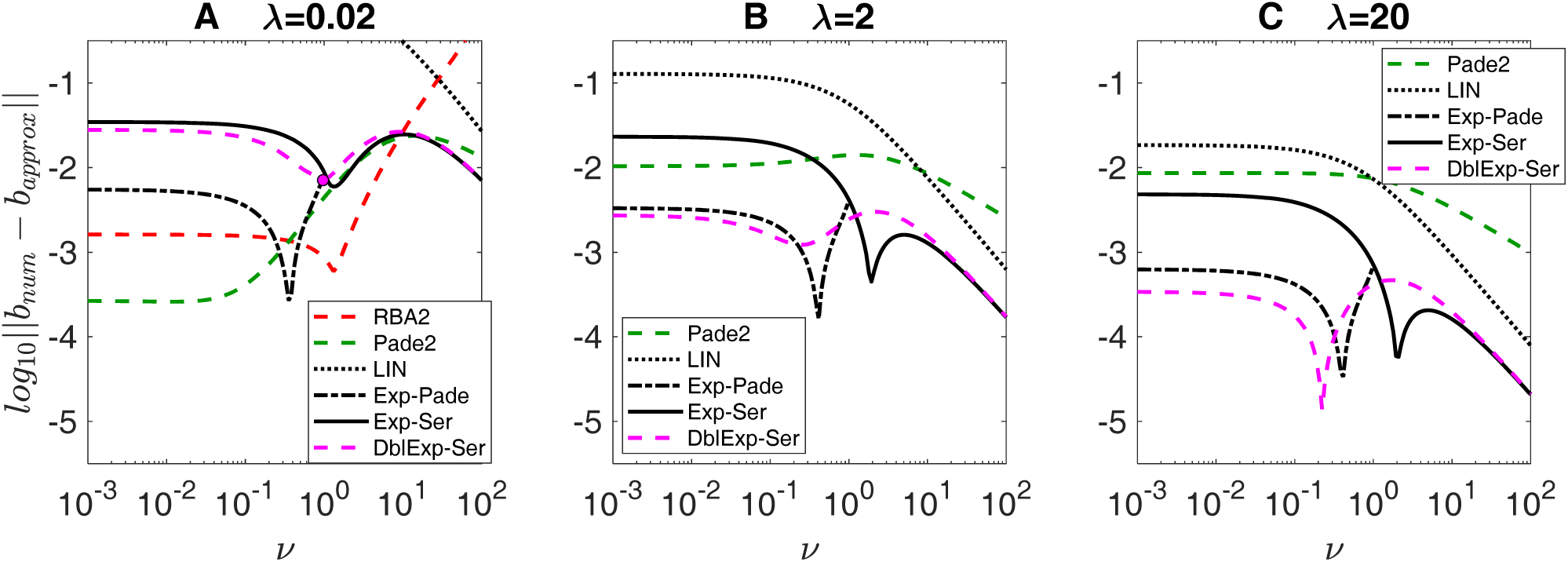
Accuracy comparison of equilibrium free buffer concentration approximations obtained by the series interpolation method: Exp-Ser (*black curves*), Exp-Padé (*dashed black curves*), DblExp-Ser (*dashed magenta curves*), and Padé2 (*dashed green curves*). Also shown is LIN (*dotted black curves*) and RBA2 (*red dashed curve*). RBA2 is only shown in A, since it requires λ<1. All curves show the error norm given by Eq. 29, on base-10 logarithmic scale, as a function of model parameter *ν* ranging from 10^−3^ to 10^2^, for 3 distinct choices of *λ*: *λ*=0.02 (A), *λ*=2 (B), and *λ*=20 (C), with *η* =1. Since Exp-Padé only yields a solution for *ν* < *η* =1, the corresponding curves terminate at *ν*=1.. *Magenta circle* in (*A*) indicates the value of *ν* below which the exponent parameter *α* of DblExp-Ser becomes imaginary (this occurs for λ<1.8). For smaller value of *ν*, the *magenta* curve in *A* corresponds to the real part of Eq. 26.

**Figure 3.**
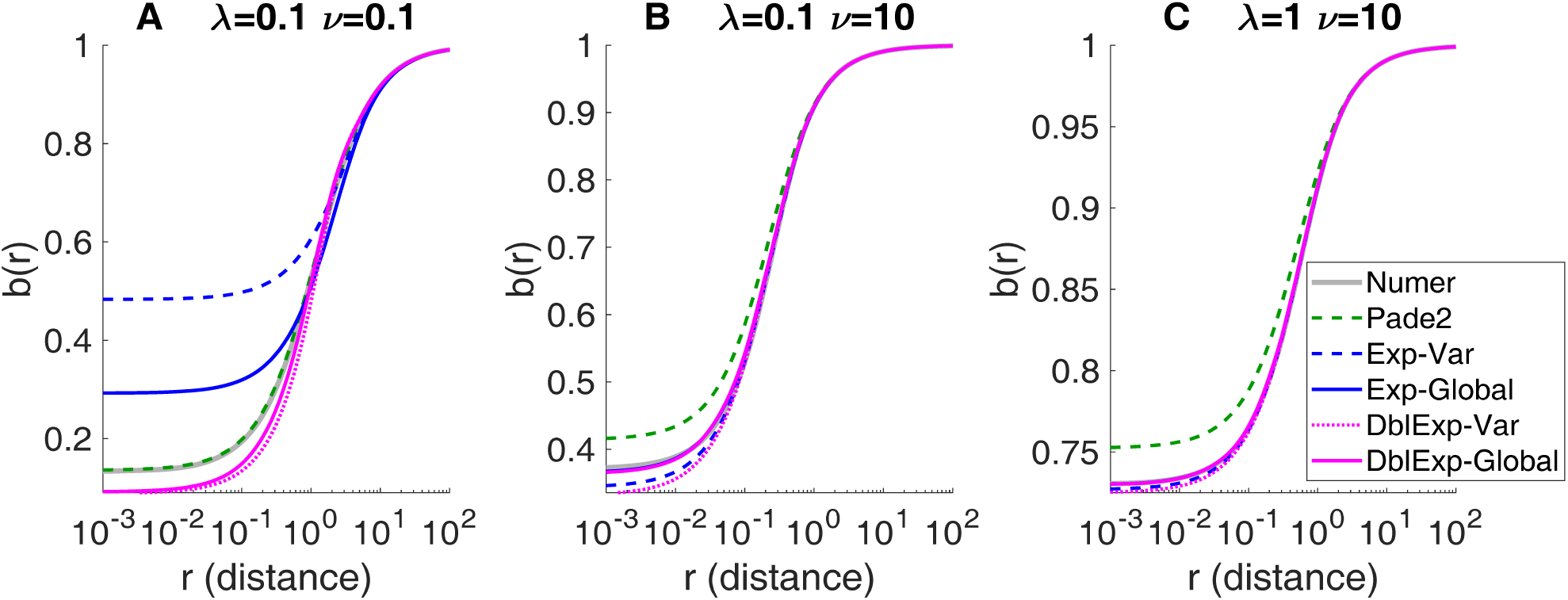
Comparison of equilibrium buffer concentration approximants obtained using the variational and the modified variational (global) methods: Exp-Var (*dashed blue curves*), DblExp-Var (*dotted magenta curves*), Exp-Global (*blue curves*), and DblExp-Global (*magenta curves*). Padé2 is also shown for comparison (*dashed green curves*). All panels show the free dimensionless buffer concentration as a function of distance from the Ca^2+^ channel, for 3 distinct choices of model parameters *λ* and *ν*, with *η*=1. *Grey curves* show the accurate numerical solution. In (A), the real part of DblExp-Var and DblExp-Global is shown. In (B) and (C), the curves for Exp-Global and DblExp-Global overlap the numerical solution.

**Figure 4.**
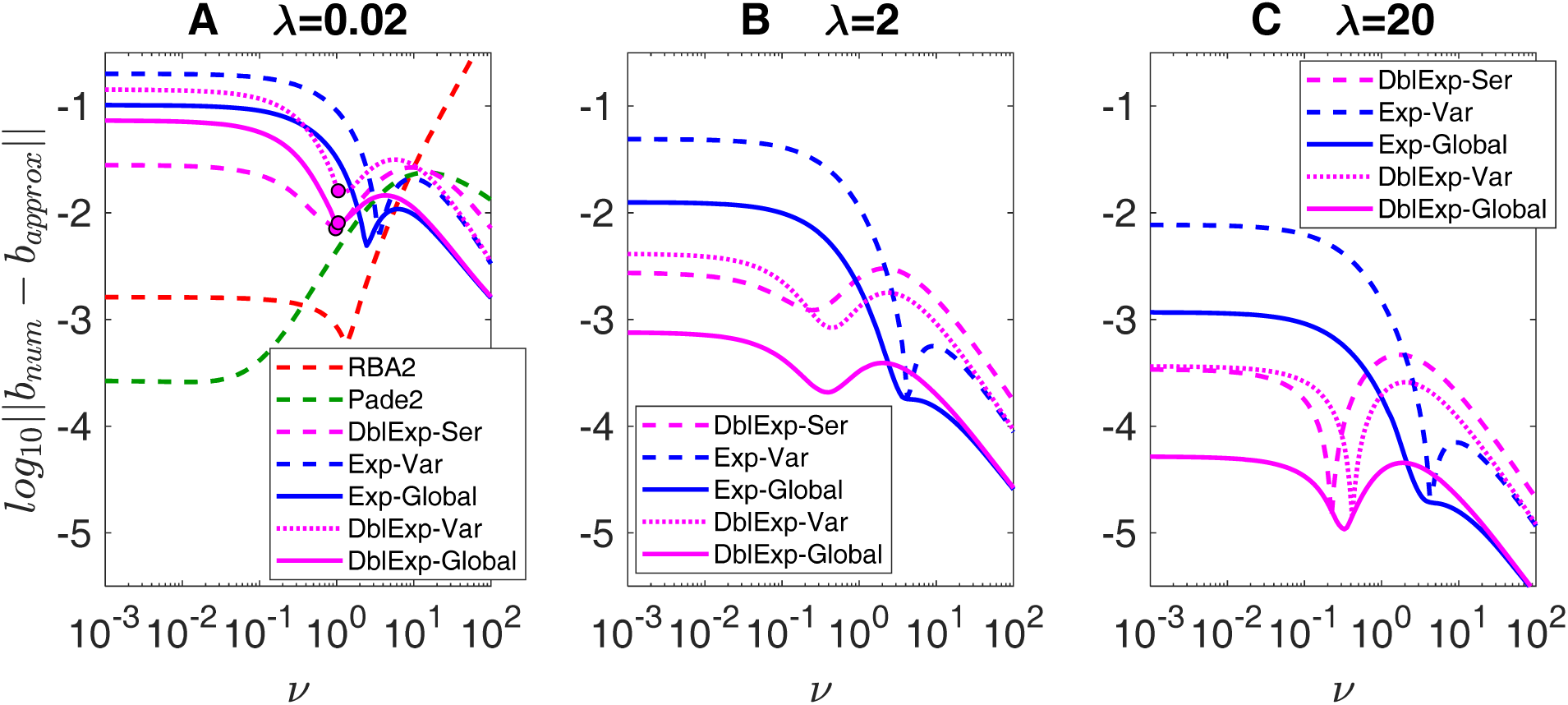
Accuracy comparison of equilibrium nanodomain free buffer concentration approximations obtained by the variational and modified variational (global) methods: Exp-Var (*dashed blue curves*), DblExp-Var (*dotted magenta curves*), Exp-Global (*blue curves*), and DblExp-Global (*magenta curves*). For comparison, also shown is the error of DblExp-Ser (*dashed magenta curves*), and (A) shows the errors of RBA2 (*dashed red curves*) and Pade2 (*dashed green curves*). All panels show the average absolute deviation of free dimensionless buffer concentration (Eq. 29), on base-10 logarithmic scale, as a function of buffer strength parameter *ν* ranging from 10^−3^ to 10^2^, for 3 distinct choices of fixed model parameter *λ*: *λ*=0.02 (A), *λ*=2 (B), and *λ*=20 (C), with *η*=1. *Magenta circles* in (*A*) mark values of *ν* below which parameter *α* becomes imaginary for the corresponding DblExp method. For these smaller value of *ν*, the magenta curves in (*A*) represent the accuracy of buffer concentration given by the real part of Eq. 26.

**Figure 5.**
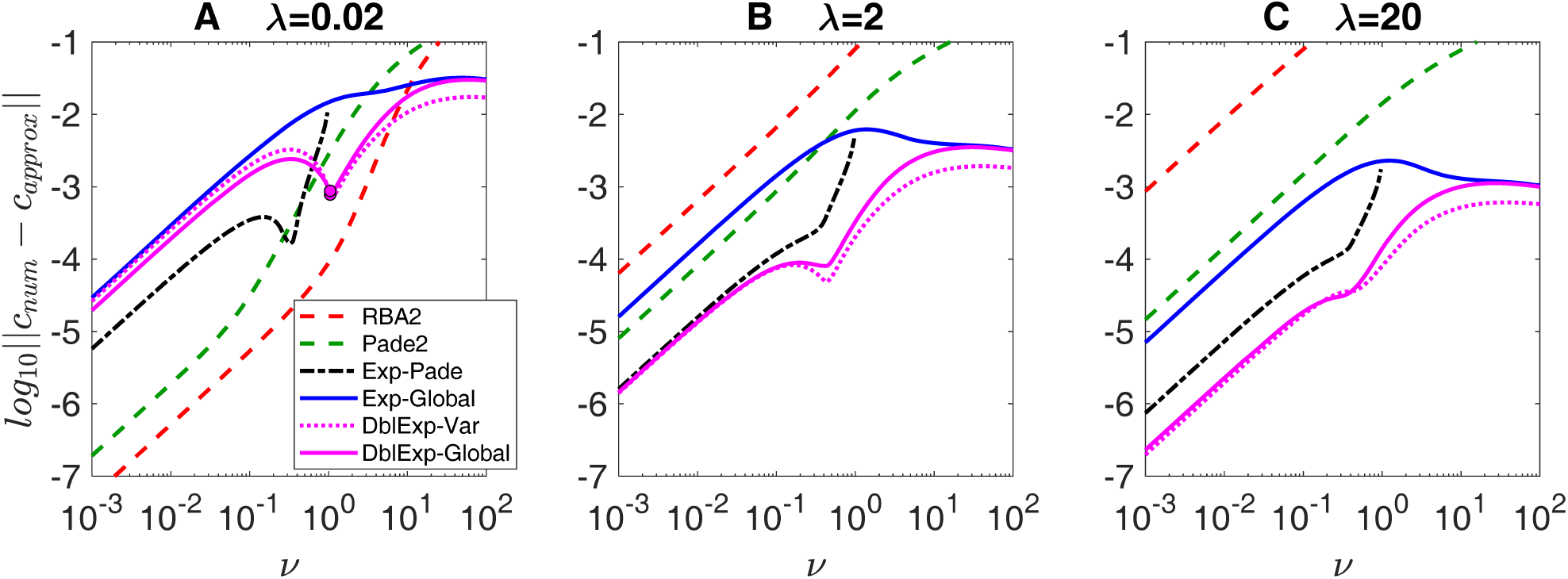
Accuracy comparison of equilibrium nanodomain Ca^2+^ concentration estimation by select optimal approximations (methods with smallest error): RBA2 (*red dashed curves*), Padé2 (*dashed green curves*), Exp-Padé (*dot-dashed black curves*), Exp-Global (*blue curves*), DblExp-Global (*magenta curve*s), and DblExp-Var (*dotted magenta curve*s). All panels show average absolute deviation of free dimensionless Ca^2+^ concentration (Eq. 36), on base-10 logarithmic scale, as a function of buffering strength parameter *ν* ranging from 10^−2^ to 10^2^, for 3 distinct choices of diffusivity parameter *λ*: *λ*=0.02 (A), *λ*=2 (B), and *λ*=20 (C), with *η*=1. Curves for Exp-Padé (*dashed black curves*) terminate at *ν*=1.

**Figure 6.**
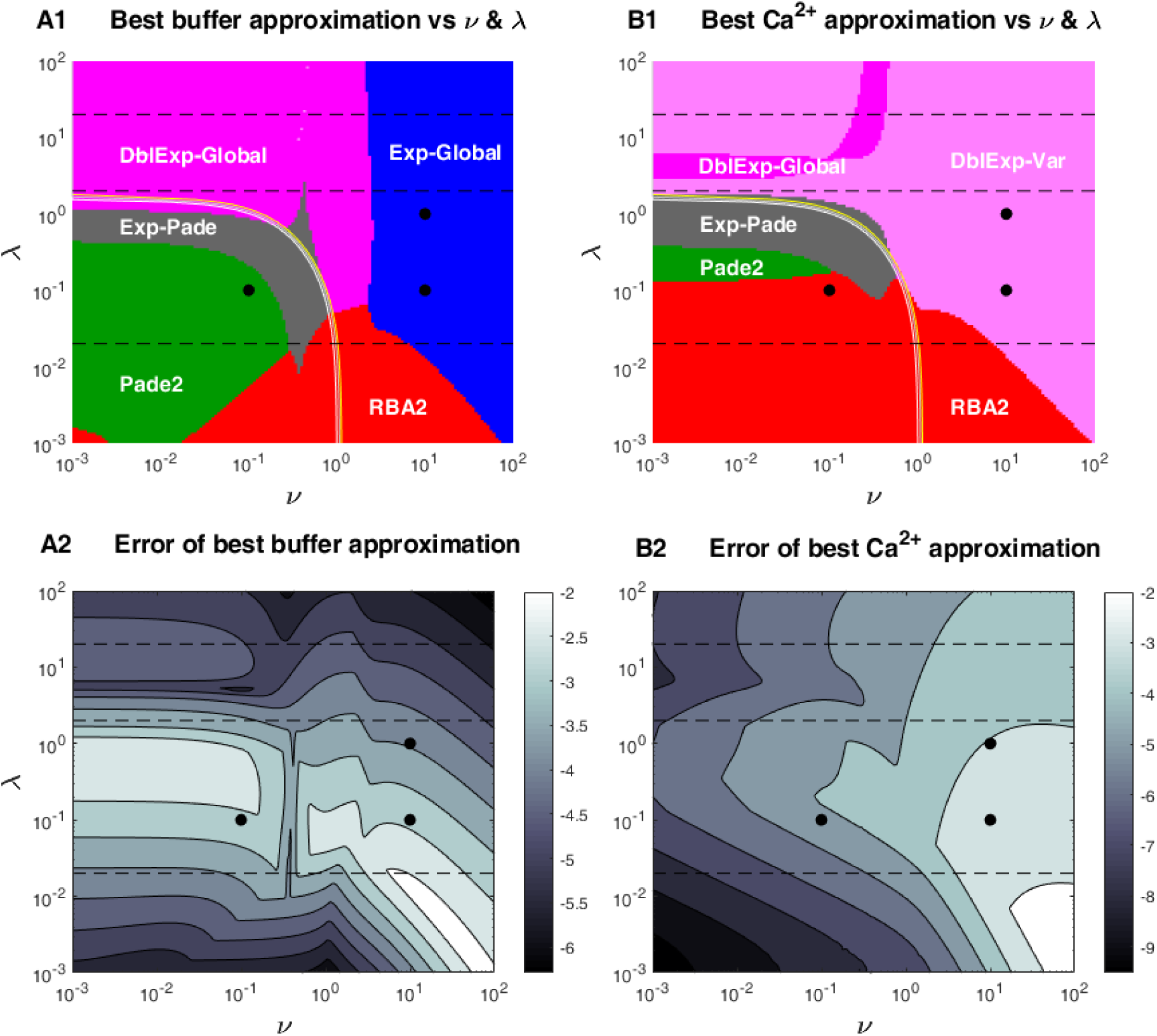
Comparison of parameter regions where a given approximant outperforms the rest in estimating (*A1*) free buffer and (*B1*) Ca^2+^ concentration in the (*ν, λ*) parameter plane, according to the error measures given by Eqs. 28 and 36. In all panels, *η*=1. Colors indicate parameter region of best performance for each approximant: Padé2 (*green*), RBA2 (*red*), Exp-Padé (*gray*), DblExp-Var (*pink*), DblExp-Global (*magenta*), Exp-Global (*blue*). Black circles corresponds to parameter values in Figs. 1,3, and dashed lines corresponds to the parameter sweep curves in Figs. 2,4-6. Thin light semi-circular curves indicate the boundaries inside of which the exponent parameters α in the DblExp-Var and DblExp-Global methods becomes imaginary (α is always real outside of the region marked by these curves, for *ν* > 1 and *λ* > 1.8). Lower panels show the smallest error in estimating buffer (A2) and Ca^2+^ (B2) concentrations achieved using the optimal approximants shown in top panels. The grayscales in A2 and B2 indicate the log-10 error values given by Eqs. 28 and 36, respectively

## III. RESULTS

### III.1 Local properties of stationary nanodomain solution

We start by generalizing some of the results previously presented in (32), without the restriction of binding-independent buffer mobility. We seek a solution to Eq. 18 which is bounded and analytic, and therefore it can be expanded in a Taylor series in *r* using the Frobenius method:

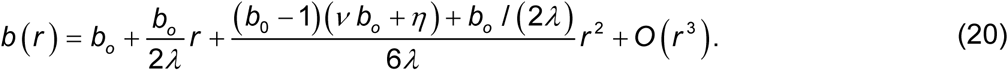

The usefulness of this series by itself is limited since the value of buffer at the channel location, *b*_o_, is *a priori* unknown, as mentioned above. Further, the convergence radius is finite due to the movable non-pole singularities of the solution in the complex *r* plane. However, the relationship between Taylor coefficients in this expansion can be used to constrain parameters of an appropriately chosen approximation. Further, by making a coordinate mapping *x* ≡ 1/ *r*, we transform our original Eq. 18 to the form:

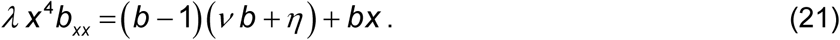

This reveals an essential singularity at *x*=0. In fact, numerical study shows that the analytic extension of *b*(*x*) to the complex-*x* plane has a branch cut across *x*=0, jumping from the physical value *b*=1 at *x*=0^+^ (*r* = +∞) to the unphysical value *b* = −*η* /*ν* at *x*=0^−^ (*r* = −∞) (see Fig. 7 in (32)).

Given that the boundary condition infinitely far from the channel is known, *b*(*x*=0^+^)=1, one can readily find the coefficients of a unique asymptotic power series expansion near x=0^+^:

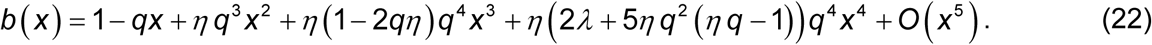

Here we used parameter *q* = 1/(*η*+*ν*) to simplify the coefficients (cf. Eqs. 16,34 in (32)). Note that terms of this long-range expansion agree up to order O(*x*^3^) with RBA and up to order O(*x*^5^) with RBA2 (Table 1), indicating that the reaction is approximately at equilibrium far from channel.

The Padé method introduced in (32) and shown in Table 1 simultaneously matches leading terms of the two expansions given by Eqs. 20 (containing unknown *b*_0_ as a free parameter) and 22, using a simple rational function interpolant, with coefficients of this rational function found as functions of model parameters *λ, ν* (or *q*), and *η*. The simplest Padé interpolant of order 1 yields:

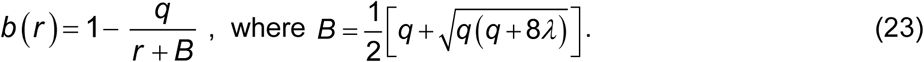

This simple function satisfies both *b*(*r*) = *b*_0_ + *b*_0_ *r* / 2*λ* + *O* (*r*^2^) and *b*(*x*) =1− *qx* + *O* (*x*^2^). The corresponding estimate of free buffer concentration at the channel location is *b*_0_ = 1 − *q* / *B*.

The Padé approximation (see Table 1) was chosen in (32) because of its algebraic simplicity and its straightforward expansion in power series in both *r* and *x*=1/*r*. Therefore, it represents an *ad hoc ansatz*, and for a fixed polynomial order, it is not necessary the most natural nor the most accurate interpolant between the short-range and long-range power series given by Eqs. 20-22. Further, although it does converge to the true solution with increasing order, closed-form expressions for its coefficients can only be obtained for the 2 lowest orders listed in Table 1. However, we observe that *all* approximants in Table 1 can be viewed as interpolants between the Taylor series in *r* and asymptotic power series in *x*=1/*r*, and therefore the series interpolation method first introduced in (32) can and should be applied to the corresponding functional forms, as well. Particularly promising in this respect is the simple exponential form of the EBA and LIN approximations, which are close to each other when *v* ≫1, and which match in this limit the first two terms in the asymptotic expansion in Eq. 22, *b*(*x*) = 1 − *qx* + *O*(*x*^2^). In fact, standard analysis by substitution *b*(*x*) = 1 − *qx* + *e*^*S*^(^*x*^) reveals that in the limit *x* = 1/*r* → 0^+^, the behavior of the general solution to Eq. 21 is described by:

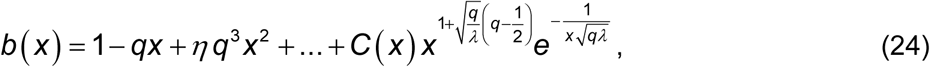

where *C*(*x*) is bounded at *x*=0. Apart from the fractional power of x, this expression has a similar form to the EBA and LIN approximations in Table 1, suggesting that the corresponding functional form is a natural *ansatz* for describing long-range behavior of the solution.

### III.2 Functional form of approximants

Given above analysis, we introduce approximants that have a simple functional form inspired by EBA and LIN, and which match the long-range asymptotic behavior of the solution, as given by Eq. 24. Namely, we consider approximations in one of the following three parametric forms:

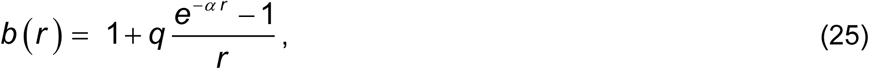

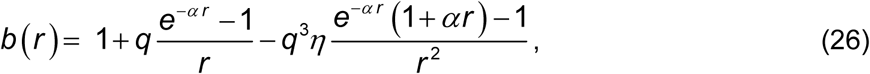

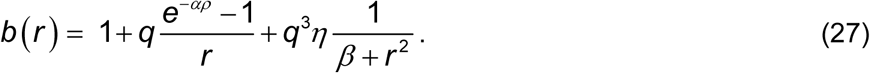

We refer to these approximants as exponential (Exp), double exponential (DblExp), and exponential-Padé (Exp-Padé), respectively. In the limit *r*→ +∞ (*x*=1/*r* → 0^+^), they explicitly satisfy the asymptotic expansion *b*(*x*) = 1 − *qx* + *η q*^3^*x*^2^ + *O*(*x*^3^) to either 1^st^ or 2^nd^ order in *x*, and are analytic at *r* = 0. The Exp and DblExp approximants depend on a single parameter *α*, while Exp-Padé contains an additional parameter *β*. Note that Eq. 25 reduces to LIN or EBA when *α* equals 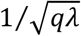 or 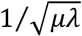, respectively (see Table 1). The novelty of our approach is that we constrain the values of parameters *α* and *β* using one of the following methods, described in detail further below:

1. Series interpolation: in this case approximants given by Eqs. 25, 26 are referred to as Exp-Ser and DblExp-Ser, respectively.
2. Variational approach: Eqs. 25, 26 in this case are referred to as Exp-Var and DblExp-Var.
3. Global method (modified variational approach): Eqs. 25, 26 will be called Exp-Global and DblExp-Global.

The value of parameter *α* is given by the solution of a quadratic equation for the exponential *ansatz*, and cubic equation for the double exponential *ansatz*, as given in Table 2. Parameters of the Exp-Padé approximant are defined by a 4^th^-order polynomial equation, and are explicitly shown in Appendix 2.

**Table 2.**
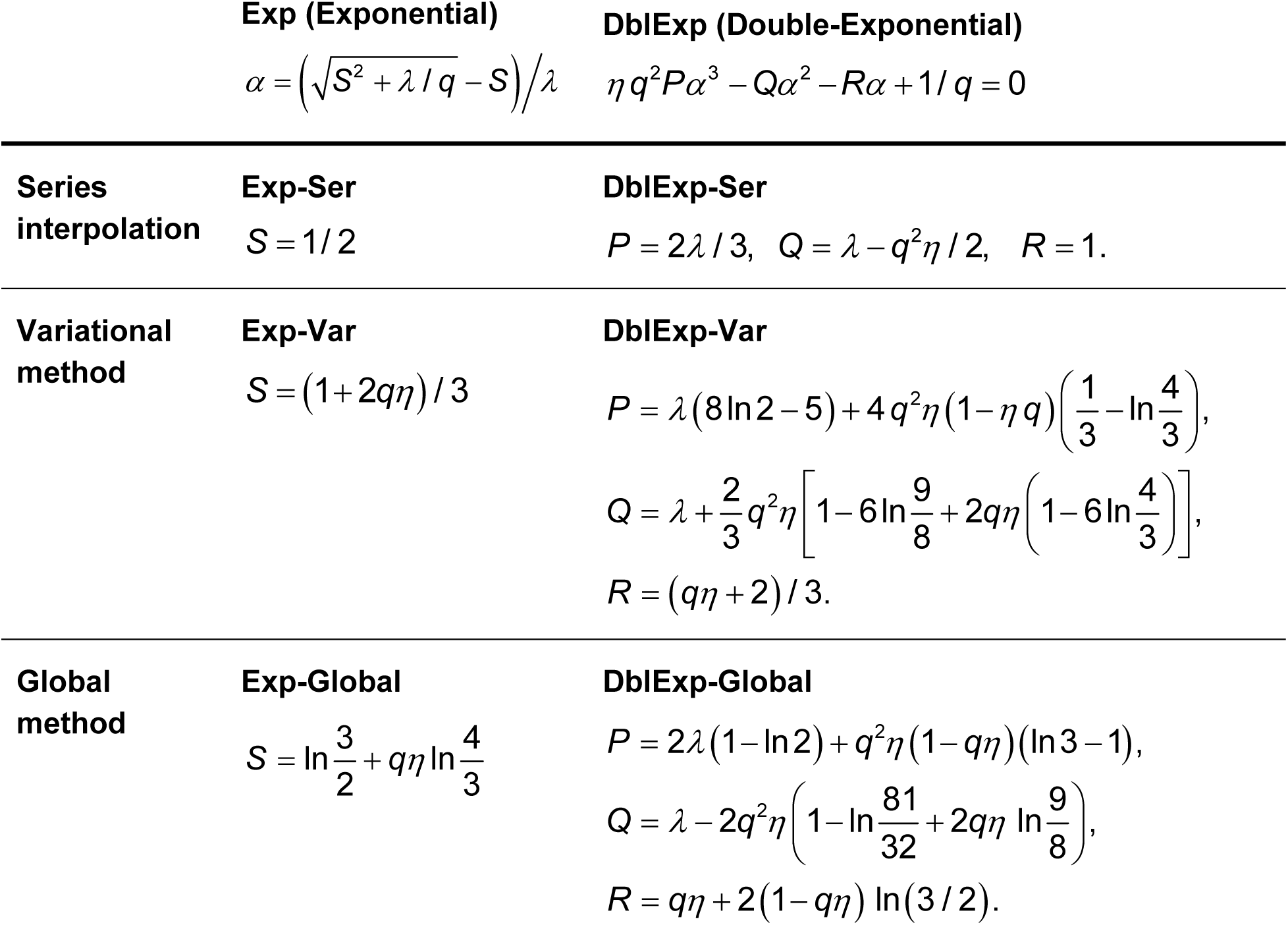
Equations for determining ansatz exponent parameter *α*. The approximants given by Eqs. 25 and 26 depend on a single constant exponent factor *α* that in turn depends on model parameters *λ, q*=(ν+*η*)^−1^ and *η* through the solution of a quadratic or a cubic equation. For all three mono-exponential approximants (Exp-Ser, Exp-Var, Exp-Global), the value of *α* is given by a solution to a quadratic equation of the same kind, but with different values of parameter *S*. Note that setting *S*=0 yields the linear approximation (LIN in Table 1). For all three double-exponential approximants (DblExp-Ser, DblExp-Var, DblExp-Global), the value of parameter α is given by a solution to a cubic equation of the same type, shown in the top row of the Table, but with different values of polynomial coefficients *P*, Q, and *R*. The positive real root of the cubic equation is given in Appendix 1.

#### III.2.1 Series interpolation approach: results

For the simple exponential *ansatz*, Eq.25, the relationship between the first two coefficients in the Taylor series in Eq. 20, *b*_l_ = *b*_0_ / 2*λ*, is satisfied for a unique value of exponent factor *α* given by a root of a quadratic equation, and listed in Table 2: 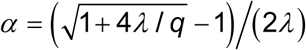. The corresponding approximant will be referred to as Exp-Ser, in contrast to LIN, which has the same functional form, but with the exponent factor value of 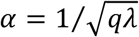 (cf. Table 1).

A slightly more complex expression in terms of two exponentials, Eq. 26, allows to match two terms in the long-range asymptotic *x*-power series, 1 − *qx* + *ηq*^3^ *x*^2^ + *O*(*x*^3^). The relationship between the first two coefficients in the Taylor series in Eq. 20, *b*_l_ = *b*_0_ / 2*λ*, holds when the value of exponent factor *α* satisfies a cubic equation given in Table 2. This cubic has at most one real positive root for all values of model parameters {*λ, q, η*}, which has an explicit solution shown in Appendix 1. The correspoding approximant will be referred to as DblExp-Ser. We note that α becomes imaginary for sufficiently small *λ* and *ν*, inside a parameter region marked by thin lines in Fig. 6A1,A2; in that case the real part of Eq. 26 will be used to compare it with other methods.

Finally, the *ansatz* given by Eq. 27 has an exponential term with parameter *α*, and a rational term with parameter *β*. Two free parameters allow to match two relationships between the first three Taylor coefficients in the short-range series expansion given by Eq. 20. This results in a polynomial system of order 4, with the level of complexity similar to that of the second-order Padé approximation (32). This polynomial system and the explicit expression for its roots are provided in Appendix 2. We note that the real positive solution for parameters *α* and *β* is only possible when ν<*η* (equivalently, 2*η q* > 1), so this approximation is not applicable for ν>*η*.

Figure 1 compares the three approximants described above (Exp-Ser, DblExp-Ser, Exp-Padé) with the previously developed Padé series interpolants of two lowest orders, as well as RBA2 (Fig. 1A), and LIN (Fig. 1C). The accurate numerical solution is shown as a gray curve. For the parameters in Fig. 1A (*λ*=*ν*=0.1), Exp-Ser (*black curve*) isn’t as accurate as other approximants, but the accuracy of Exp-Padé (*dashed black curve*) and DblExp-Ser (*dashed magenta curve*) is excellent, and comparable to that of Padé2 (*dashed green curve*); in fact, the three curves completely overlap with the numerical solution curve. This is despite the fact that *α* in DblExp-Ser expression is complex for *λ*=*ν*=0.1, so this is not an optimal parameter region for DblExp-Ser, and the real part of Eq. 26 is used in this case. For larger values of *λ* and/or *ν* in Figs. 2*B* (*λ*=0.1, *ν*=10) and 2*C* (*λ*=1, *ν*=10), approximants Exp-Ser and DblExp-Ser are more accurate than Padé and even Padé2. These results suggest that these series interpolants may be superior to previously developed approximants in estimating Ca^2+^ nanodomains in a wide range of model parameters. Among previously developed approximants listed in Table 1, only RBA2 provides comparable accuracy, in the case λ<1, corresponding to parameters in Fig. 1A (*dashed red curve*).

Comparing the results by eye for several combinations of model parameters is clearly insufficient to unveil the parameter-sensitivity of approximant accuracy; in fact, the difference between several approximants is almost impossible to tell from Fig. 1. Therefore, following prior work (20,31,32), we explore parameter dependence of the absolute deviation between the given approximation *b*_approx_ and the accurate numerical solution, *b*_numer_:

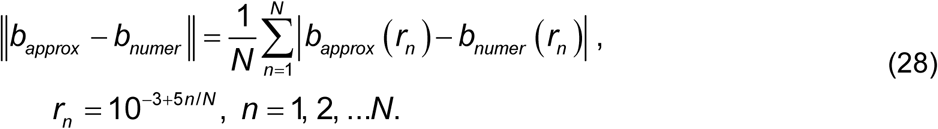

The deviations are computed on a set of *N*=100 points spanning 5 orders of magnitude of distance *r*, from 10^−3^ to 10^2^. Since we use logarithmically spaced points, this norm is equivalent to an *L*^1^ norm weighted by 1/*r*, and therefore it requires a short-range cut-off (we pick *r* ≥ 10^−3^). The higher weight at small *r* is justified by the fact that the short distance range is of greater interest, physically. Fig. 1 indicates that the chosen range of *r* is sufficient to capture the qualitative behavior of solutions for a wide range of parameter values. We checked that none of the conclusions are changed qualitatively by choosing an *L*^∞^ norm instead.

The parameter dependence of this error norm is shown in Fig. 2, as the value of *ν* is systematically varied from 10^−2^ to 10^2^, for three distinct values of *λ*. Each curve shows the error measure given by Eq. 28 for the corresponding approximation. For the sake of comparison, also shown are the error of the 2^nd^ order Padé interpolant (Padé2, *dashed green curves*), the linear approximant (LIN, *dashed black curves*), and RBA2 (*dashed red curve*, Fig 2A only). For smaller values of *λ* (Fig. 2A), Padé2 and RBA2 are still the superior approximation methods, but with increasing *λ*, the exponential series interpolation approximants outperform all approximants in Table 1 in estimating free buffer concentration. Thus, the choice of the optimal approximation method depends on the particular combination of model parameter values.

### III.3 Variational approach

We now consider a completely different method of approximating solutions, based on a variational approach. As we rigorously demonstrate in Appendix 3, the solution to Eq. 14 represents a unique minimizer of the following functional, in an appropriate function space:

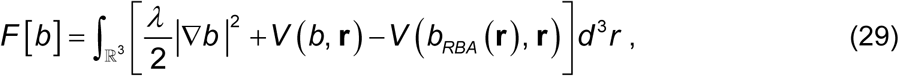

where *V*(*b*, **r**) is defined by

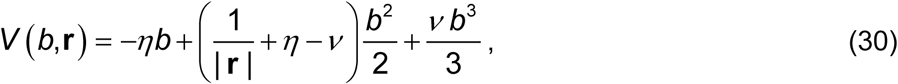

and *b*_RBA_(**r**) is the 1^st^-order RBA approximants given in Table 1, which solves Eq. 14 when *λ*=0. Subtraction of *V*(*b*_RBA_(**r**), **r**) in Eq. 29 is necessary to ensure boundedness of *F*[*b*]. Considering perturbations *b* → *b* + *εϕ*, where *ϕ* is a smooth function with compact support 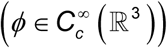, and denoting *V* ‘(*b*, **r**) the 1^st^ partial derivatives with respect to *b*, the variational derivative (the Gâteaux derivative) of *F*[*b*] in the direction of *ϕ* is

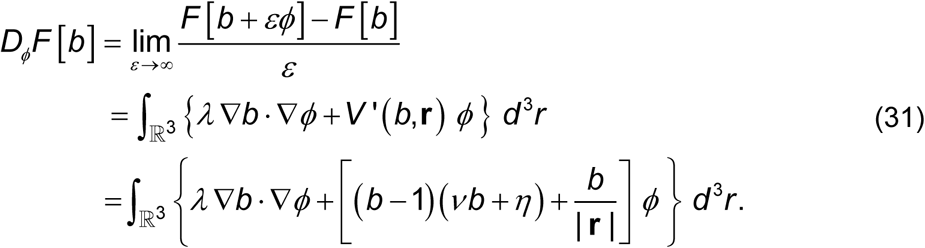

Therefore, setting

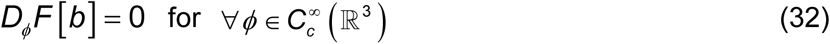

formally yields the weak form [42] of Eq. 14. As is proved in Appendix 3, the minimizer of *F*[*b*] is unique and radially symmetric. Therefore, we seek an *ansatz* of the form *b*(*r*; *α*_*k*_), and consider perturbations with respect to the *ansatz* parameters *α*_*k*_, i.e. we take *ϕ* = ∂*b* (*r* ; *α*_*k*_) / ∂*α*_*k*_. Performing integration by parts in the derivative term transforms Eqs. 31-32 to

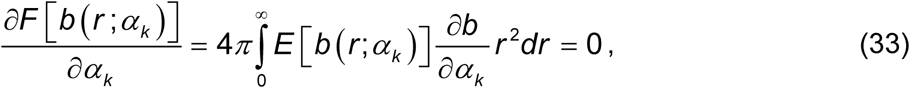

where *E*[*b*] is defined in Eq.18. For the *ansatz* given by Eqs. 25-27, this integral may be computed in closed form, allowing to obtain the optimal values of parameters *α*_*k*_ by differentiation. For the lowest-order exponential *ansatz* (Eq. 25), considering *b*(*r* ;*α*) with one free parameter in Eq. 33 leads to a quadratic equation for *α* with a unique positive real root, as given in Table 2. The corresponding approximant will be referred to as Exp-Var (see Table 2). Note the similarity in the expression for *α*, as compared to the series interpolant method result in Table 2.

For the more complicated case of a double exponential *ansatz* (Eq. 26), Eq. 33 leads to a cubic rather than a quadratic equation for *α*, analogously to the series interpolation method; this cubic is shown in Table 2, and its closed-form solution is given in Appendix 1. This cubic has a single real positive real root for a wide range of model parameters {*λ, ν, η*}, and we refer to the corresponding approximant as DblExp-Var. However, just like in the case of DblExp-Ser, *α* becomes complex when both *λ* and *ν* are sufficiently small. In this parameter regime, the real part of Eq. 26 still provides an accurate approximant. The performance of Exp-Var and DblExp-Var approximants will be investigated below, after considering our final approximation method.

### III.4 Global method: modification of the variational approach

Given that Eqs. 25-26 represent narrow classes of functions that cannot provide a true minimum of *F*[*b*], it may be useful to consider modifications of Eq. 33 that allow to achieve a lower value of our chosen error norm given by Eq. 28. One such modification is to replace the Jacobian factor *r*^2^ in Eq. 33 with the first power of *r*, increasing the contribution of small distances in this integral, and thereby potentially reducing the error at short range:

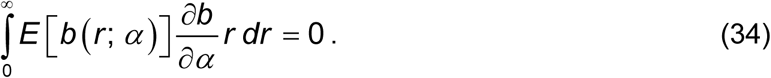

We refer to this method of setting approximant parameter values as the *Global* method, or modified variational method. Eq. 34 can be rigorously obtained from the variational derivative given by Eqs. 31-32, but this time applied to perturbations *ϕ* of form

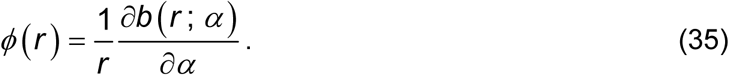

We note that for the *ansatz* in Eqs. 25-26, this perturbation remains finite as *r*→0. Numerical results show that this modification *does* lead to noticeable improvement of the resulting approximants close to the channel location, for some combinations of model parameters. In fact, for some parameter regimes this method clearly outperforms the series interpolation and the variational approaches with respect to the weighted *L*_1_ error measure given by Eq. 28.

For the lowest order exponential *ansatz* (Eq. 25), after replacing *b*(*r*; *α*) in Eq. 34 with Eq. 25, one obtains a quadratic equation for *α* with a single positive real root given in Table 2; we refer to the corresponding approximant as Exp-Global. Just as in the case of the series intepolant method and the variational method, applying this method to the double exponential *ansatz* (Eq. 26) leads to a cubic equation for parameter α, given in Table 2. We verified that this cubic has a single real positive real root for a wide range of model parameters {*λ, ν, η*}, and we refer to the corresponding approximant as DblExp-Global. However, like in the case of DblExp-Ser and DblExp-Var approximants, parameter *α* becomes imaginary when both *λ* and *ν* are sufficiently small; in that case, the real part of Eq. 26 will be used.

We note that a more straightforward approach of minimizing a weighted *L*^2^ norm of *E*[*b*] also leads to a closed-form solution in the case of a single-exponential *ansatz*, but the resulting approximant does not perform significantly better than the ones we present above, and its parameter *α* is given by solution to a more complicated 4^th^ order polynomial equation.

### III.5 Accuracy of the variational and global approximants

Figure 3 compares all variational and global approximants described above (Exp-Var, DblExp-Var, Exp-Global and DblExp-Global) with Padé2 and the accurate numerical solution, using the same combination of model parameters as in Fig. 1. It shows that in some cases (Fig. 3B,C) the global approximations are more accurate than Padé2 and other series interpolants (cf. Fig. 2B,C). Further, in Figs. 3B and 3C, global approximants perform better than the corresponding variational approximants, and the differences between global methods and numerical results are barely noticeable. In contrast, Fig. 3A illustrates that for *ν*=*λ*=0.1, none of the variational and global approximants are as accurate than Padé2, suggesting that the series interpolation methods may be superior for small values of *ν* and *λ*. We conclude the variational method and the global method can be great improvements compared with the series interpolation method in some, but not all, parameter regimes.

Figure 4 shows a more systematic comparison to reveal the accuracy of the approximants obtained using the variational and the global methods in more detail. As in Fig. 2, the value of *ν* is systematically varied from 10^−2^ to 10^2^, for three different fixed values of dimensionless buffer diffusivity parameter *λ*. Each curve shows the average absolute error in buffer concentration approximation, as given by Eq. 28. The error of the series interpolant DblExp-Ser is also shown for comparison in all panels, while Fig. 4A also shows the accuracy of RBA2 and Pade2. For small values of *ν* and *λ* (Fig. 4A), RBA2, Pade2, and even DblExp-Ser are outperforming the global approximants. However, as one increases the values of *ν* and *λ*, global approaches are starting to show their advantage. For most parameter regimes, approximations obtained using the modified variational (i.e. *global*) method are more accurate than the corresponding approximations obtained using the unmodified variational method. For example, in all panels of Fig. 4, Exp-Global (*blue curves*) is superior to Exp-Var (*dashed blue curves*).

We note that the 2nd term in the DblExp approximants reflects the 2nd term in the long-range asymptotic series, which scales as *q*^3^=1/(*η*+ν)^3^, therefore the double-exponential and the mono-exponential *ansatzes* become equivalent when *q* is sufficiently small, corresponding to large values of buffer strength parameter ν. This behavior of accuracy as *ν*→∞ is apparent in Figs. 2 and 4.

As noted above, Ca^2+^ concentration is uniquely determined from the equilibrium buffer concentration through the Ca conservation law, Eq. 11. Nevertheless, it is useful to look separately at the accuracy of the Ca^2+^ estimation by the methods we present. Close to the channel location Ca^2+^ concentration is dominated by the unbounded point source term, 1/*r*, and therefore we will use a logarithmic error measure when comparing Ca^2+^ concentration approximations (20,31,32):

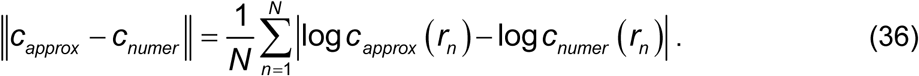

This sum extends over the same logarithmically spaced points that were used for the buffer error measure given by Eq. 28, namely a set of 100 points spanning 5 orders of magnitude of distance.

Figure 5 plots this Ca^2+^ error measure for the optimal approximations that achieve the greatest accuracy over the wide range of model parameters *λ* and *ν*. Because of the difference between the buffer and the Ca^2+^ error measures (cf. Eq. 28 vs. Eq. 36), the accuracy profile of different Ca^2+^ concentration approximants shown in Fig. 5 doesn’t match perfectly the accuracy of the corresponding free buffer approximants shown in Figs. 2 & 4, despite the one-to-one relationship between the Ca^2+^ concentration and free buffer. As explained above, the relative error in Ca^2+^ concentration estimation is particularly sensitive to the accuracy of the method at intermediate distances, rather than its accuracy in the vicinity of the channel, as is the case for the free buffer error measure (20,32). Note in particular that the DblExp-Var or DblExp-Global yield the most accurate Ca^2+^ approximations for *λ*≥1 (see Figs. 5B,C), contrary to the error in buffer estimation, which is minimized by the Exp-Global and DblExp-Global approximants (cf. Fig. 4B,C). However, for small values of *λ*, RBA2 and Pade2 are the best methods for estimating both Ca^2+^ and buffer concentration (Figs. 4A, 5A).

Finally, Fig. 6 summarizes all results presented in Figs. 1-5, marking the best approximants and their errors for a wide range of buffer mobility *λ* and buffering strength *ν* varying over 5 orders of magnitude. It shows that the methods we presented significantly improve the accuracy of approximation for a wide range of model parameters, and especially those corresponding to larger values of buffer mobility *λ* and buffering strength *ν*. In fact, these methods outperform all previously developed approximants with the exception of the 2^nd^-order RBA and 2^nd^ order Padé (20,32), the two methods which are still superior in particular regions of parameter space corresponding to small buffer mobility *λ* and small-to-moderate buffering strength *ν*.

## IV. Discussion

We have presented a significant extension of prior modeling work on equilibrium single-channel Ca^2+^ nanodomains, presenting two distinct approaches applied to several types of parametric approximants, which to our knowledge have not been considered previously. In particular, we extend the series interpolation methods recently used to construct rational function (Padé) approximants (32), generalizing it to more accurate and natural parametric forms given by Eqs. 25-27, which bear resemblance to the EBA and LIN approximants obtained previously using different methods. Furthermore, we also applied two modifications of the variational approach to approximants of the same functional form, resulting in significant improvement of approximation accuracy for a wide range of parameters. As summarized in Fig. 6, a combination of several of the best-performing approximants can achieve an excellent estimation for the free buffer and Ca^2+^ concentration near an open channel, for several orders of magnitude of dimensionless parameters *λ* and *ν*. In fact, Figure 6 can be used to write a simple algorithm for the selection of the optimal method. Moreover, this algorithm can be further simplified by using just three methods, Pade2, RBA2 and DblExp-Global, allowing to achieve an accuracy of 1% or better in the entire parameter range explored in Fig. 6. As Fig. 6A2,B2 shows, the parameter region posing the greatest challenge corresponds to *λ*<<1, *ν* >>1. However, Figs. 1B, 2A, 3B, 4A & 5A demonstrate that reasonable accuracy is achieved even in this parameter regime.

Although all accuracy comparisons shown in Figs. 1-6 were performed in the special case *η*=1, corresponding to zero background Ca^2+^ concentration (*c*_∞_=0) and binding-independent buffer mobility (*δ*^*^_B_=1), this was done solely for the sake of simplicity, and we explicitly verified that the approximant accuracy is not overly sensitive to the value of *η.*

We note that the accuracy profiles shown in the Figs. 2, 4-7 depend on our choice of the error measures, given by Eqs. 28, 36. For instance, without spacing mesh points logarithmically in these error measures, the accuracy ranking of different methods may change. However, this error measure choice provides a very demanding and restrictive comparison, covering a very wide range of distances, and weighting the error more at short distance from the channel (20,31,32). Therefore, we believe that the chosen error measures are appropriate and yield the best comparison method given the wide range of parameters we consider. Further, we checked that the conclusions are not substantially changed if the L^∞^ norm is chosen instead.

The drawback of the methods we present is that the expression for approximant parameters can be quite complex, especially for the *ansatzes* with more than one exponential term. The level of complexity of different methods is not the same: the simplest ones are the mono-exponential approximants (Exp-Ser, Exp-Var, Exp-Global), followed by double-exponential methods that require finding a root of a cubic equation (DblExp-Ser, DblExp-Var, DblExp-Global), and finally, two methods, Exp-Padé and Padé2, require solving a fourth-order polynomial system. However, all approximants were determined as closed-form expressions that only take several lines of computer code (see Appendices 1 and 2).

Several other functional forms not shown in Table 2 were also considered, but are not presented here since they either did not result in better accuracy compared to other approximants, or provide only a minor improvement in limited regions of parameter space while complicating the expressions for parameters. This is true for example for the double-exponential approximation given by Eq. 25 but with two different exponent parameters, *α*_l_ and *α*_2_. However, it is possible that we missed other accurate approximants. It is possible that such improved *ansatzes* could be found, for instance by taking into account the singularities of the analytic extension of the buffer concentration to the unphysical complex-distance plane. We note that only RBA captures the branch cut of this analytic extension, which jumps from the physical value *b*=1 at *x*=0^+^ (*r=+*∞) to the unphysical value *b* = −*η* / ν at *x*=0^−^ (*ρ=−*∞) (see Fig. 6 in (32)). Further, as noted above, 2^nd^-order RBA derived in (20) agrees with the long-range asymptotic expansion of the true solution given by Eq. 22 up to terms of order *x*^5^ (20,32). Therefore, our initial efforts to construct an improved *ansatz* were based on modifying the RBA approximant. However, so far we failed to find a successful modification of RBA that improves its performance.

More importantly, the presented approaches can be extended to the study of complex buffers with more realistic Ca^2+^ binding properties. Most of prior modeling efforts, including this study, focused on a simple buffer with one-to-one Ca^2+^ binding, but most biological buffers possess several binding sites with distinct Ca^2+^ binding characteristics, such as calretinin and calmodulin (39-41). To date, only RBA has been extended to such buffers, and only to 1^st^ order (31). However, our preliminary exploration reveals that the series interpolation approach can be extended to such buffers, using a combination of rational and exponential functions, which is a subject of our current work. Another direction of potential improvement is relaxing some of the key simplifying assumptions of the model, allowing for simple volumetric Ca^2+^ extrusion and extension to Ca^2+^ channel pore of a finite width (16), and exploring the generalization of these methods to the case of two or more channels.

## Author Contributions

V.M. conceived and designed the research project, V.M. and C.M. developed analytic methods, Y.C. and V.M. performed model analysis, numerical coding, numerical simulations and simulation data analysis. All Authors took part in the writing and review of the manuscript.

## Acknowledgements

This work was supported in part by NSF grant DMS-1517085 to V.M., and by NSF grant DMS-1908709 to C.M. The authors acknowledge valuable discussions with Vitaly Moroz.

## APPENDIX 1: Exponent parameter for double exponential approximations

For each of the three approximation methods summarized in Table 2, the parameter α of the double-exponential *ansatz* satisfies a cubic equation of form:

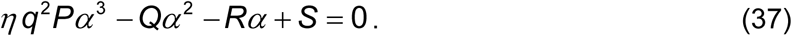

The three roots of this cubic can be succintly represented in the folowing form:

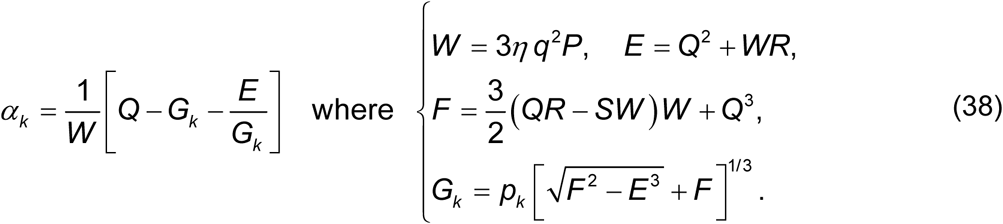

The constants *p*_*k*_ (*k* = 1,2,3) in the expression for the intermediate quantity *G*_*k*_ denote branches of (−1)^1/3^ :

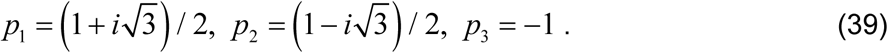

In this notation, the real positive root of Eq. 37 corresponds to the value *α*_l_ when implemented verbatim in MATLAB (Mathworks, Inc). For each of the three double-exponential approximants, the imaginary part of the root becomes non-zero for small values of ν and λ corresponding to the inner region marked by thin curves in Fig. 6A1,B1.

## APPENDIX 2: parameters of the Exp-Padé approximation

For the Exp-Pade *ansatz* (Eq. 27), matching the relationship between the first three terms in the Taylor series of the solution (Eq. 20) leads to the following 4^th^ order polynomial system for the Exp-Pade parameters *α* and *β*:

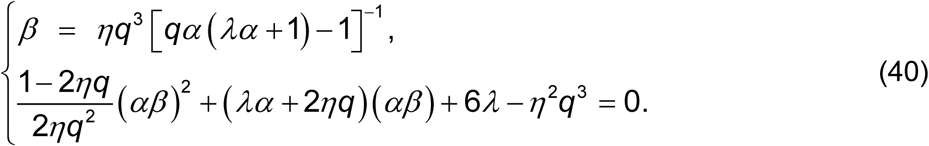

This is a fourth-order polynomial equation for *α*, with the following explicit solution:

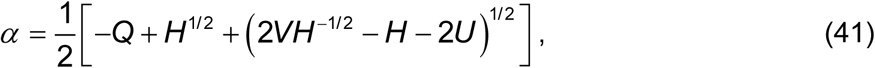

where constants *U, V, H, Q* are determined by model parameters {*λ, q, η* } according to

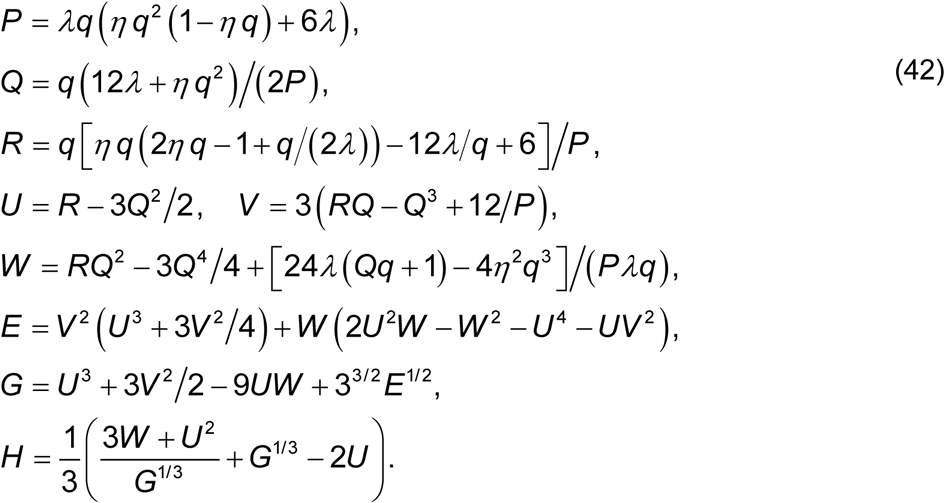

We note that the other three roots do not yield real positive values of *α* and *β*. In the case λ<<1 and ν<<1, these expressions may need conditioning to prevent numerical loss of significance.

## APPENDIX 3: Existence and uniqueness of solution

Here we outline a rigorous mathematical study of Eq. 14 to establish the basic qualitative characteristics of its biophysically relevant solutions. The solutions of this equation must be understood in the distributional sense in ℝ^3^ [42], in view of the fact that the right-hand side of Eq. 14 blows up at the origin and, therefore, the derivatives of *b*(**r**) are undefined classically at **r**=0. We will take advantage of the variational formulation, Eqs. 29-32, to establish basic existence, uniqueness, regularity and symmetry properties of the solutions of the above equation in the physically relevant class of functions *b* : ℝ^3^ → ℝ, namely functions that approach the limit at infinity sufficiently fast and obey the bounds in Eq. 16. To make the statement in Eqs. 29-32 more precise, we need to ensure that *F*[*b*] is well defined and differentiable for a given *b*. A natural class of functions ensuring these conditions is the homogeneous Sobolev space 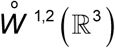, i.e., the space of locally integrable functions with square integrable first weak derivatives [43]. This makes the first term in the integrand in Eq. 29 well-defined. Nonetheless, we still need to make sure that the rest of the integrand does not give rise to a divergent integral due to a possible slow decay of *b*(**r**) − 1 as **r** → +∞. To control the latter issue, we invoke Eq. 16.

For simplicity of notation, let *V*’(*b*, **r**) and *V*’’(*b*, **r**) denote the 1^st^ and the 2^nd^ partial derivatives with respect to *b*. Taylor expanding around *b*_RBA_(**r**) and taking into account that *V*’(*b*_*RBA*_, **r**) = 0, we have

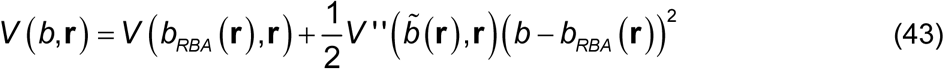

for some 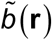 lying between *b*(**r**) and *b*_RBA_(**r**). We note that *b*_RBA_(**r**) satisfies the bounds in Eq. 16, obeys *b*_RBA_(**r**) ∼ |**r**| as |**r**|→0, and agrees up to order O(|**r**|^−3^) with Eq. 22 as |**r**|→+∞. Since 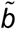 also satisfies the bounds in Eq. 16 and because

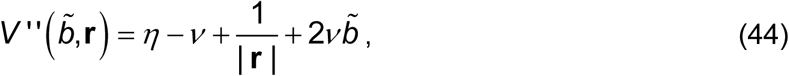

we obtain from Eq. 44 and the definition of *η* (Eq. 13) that

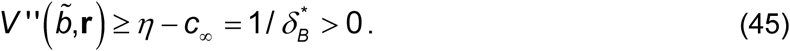

In particular, *F* is non-negative in the considered class. Also, by inspection 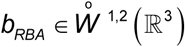. Therefore, it holds that

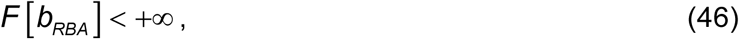

indicating that *F* is finite on a non-empty subset of 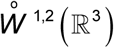 satisfying the bounds in Eq. 16.

We now proceed with establishing existence of solutions of Eq. 14 which are minimizers of *F* among all 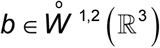 satisfying Eq. 16. To this aim, we first redefine *F* to relax the constraints in Eq. 16, by introducing

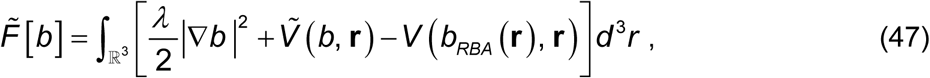

where

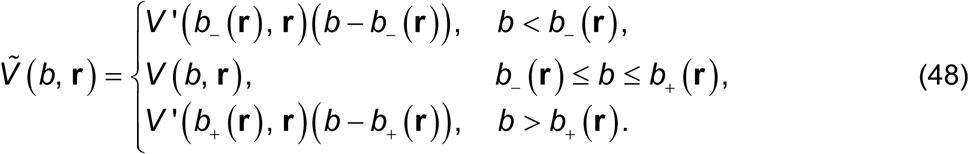

Notice that 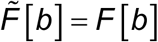 for all *b* satisfying Eq. 16. Also, by inspection 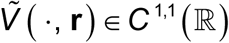 for all **r**≠0, and

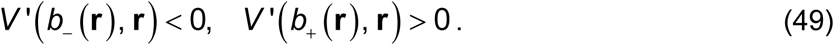

In particular, 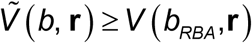 for all *b* ∈ ℝ and **r** ∈ ℝ^3^.

Next we use the direct method of calculus of variations [44] to establish existence of minimizers of 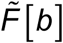. In view of Eq. 46, we have inf 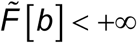. Existence of minimizers then follows from coercivity and lower semicontinuity of 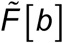 with respect to the weak convergence in 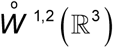 and strong convergence in 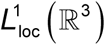 [44]. Indeed, if 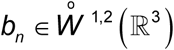 is a minimizing sequence, then for any *R* > 0 we have

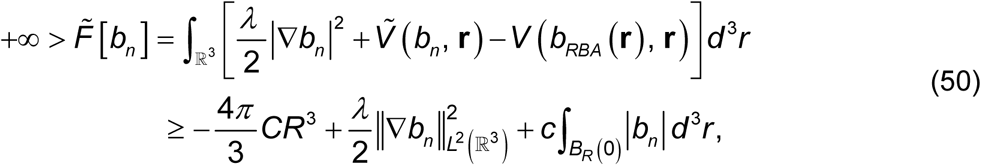

because by construction 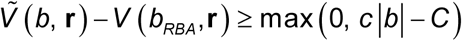 for some *c, C* > 0 and any 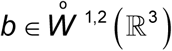 From Eq. 50 we obtain 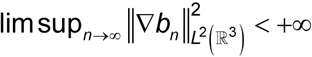 and after extraction of a subsequence we have ∇*b*_*n*_ ⇀ ∇*b* in *L*^2^ (ℝ^3^; ℝ^3^) and *b*_*n*_ (**r**) → *b* (**r**) for almost every **r** ∈ ℝ,^3^ for some 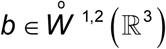. Then by lower semincontinuity of the norm and Fatou’s lemma applied to *V* (*b*_*n*_ (**r**), **r**) we get that lim 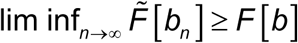, and so *b* is a minimizer of 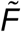. Furthermore, since 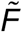 is Fréchet differentiable with respect to compactly supported perturbations, we also have 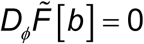, i.e.,

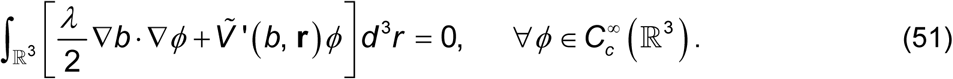

Having established existence of a minimizer of 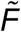, we now show that it satisfies Eq. 16 *a posteriori*. To show that *b* ≤ *b*_+_ we define 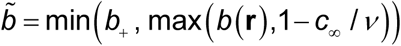; by Eq. 49 we have 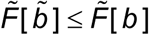, and this inequality is strict unless 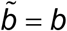 almost everywhere in ℝ^3^. Similarly, to establish *b* ≥ *b*_−_, we define 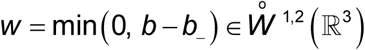, and note that *w* = 0 in *B*_1 /*ν*_ (0) or whenever *b* ≥ *b*_−_ in 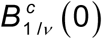. Defining 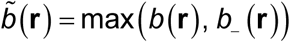, we have

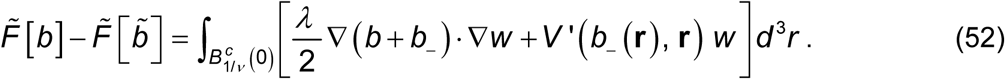

Using Eqs. 49 and 51, and the fact that 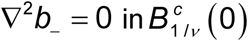 distributionally, integrating by parts we obtain

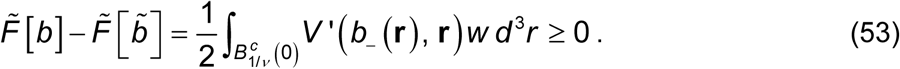

This inequality is strict unless *b* = *b*_−_ almost everywhere in ℝ^3^. Thus, the minimizer *b* satisfies Eq. 16 and, hence, is also a minimizer of *F*[*b*] in 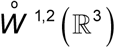, subject to the constraint in Eq. 16.

We now establish uniqueness, regularity and radial symmetry of the minimizer *b*. By Eq. 51, *b* satisfies Eq. 32 and is unique in this class due to strict convexity of *F* ensured by Eq. 45. Namely, if *b* is a minimizer and 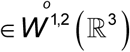 is such that *b* + *w* still satisfies Eq. 16, with the help of Eqs. 51 and 43 we can write

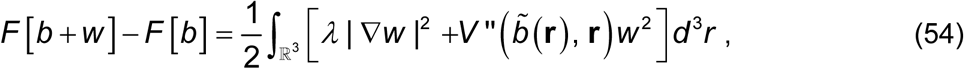

for some 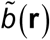 between *b*(**r**) and *b*(**r**)+*w*(**r**). So, by Eqs. 54 and 45, we have *F*[*b*+*w*] > *F*[*b*] for every *w*(**r**) ≠0, and, therefore, *b*(**r**)+*w*(**r**) is not as minimizer unless *w*(**r**)=0 almost everywhere in ℝ^3^.

Then, by uniqueness of minimizer, we have *b*(**r**)=*b*(|**r**|) (with a slight abuse of notation), i.e. *b* is radially symmetric, as minimization may be carried out in the class of radial functions to obtain a radial solution of Eq. 32. Finally, elliptic regularity theory [43] yields that for any 1 ≤ *p* < 3,we have 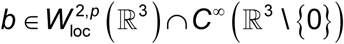, and hence by Sobolev embedding [42] we have *b* ∈ *C*^0,*α*^ (ℝ^3^) for any *α* ∈ (0, 1). In particular, *b*(*r*) is continuous at *r*=0 and solves Eq. 18 for each *r* > 0. Integrating this equation once near the origin yields

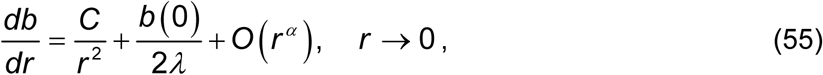

for some *C* ∈ ℝ. In view of square intrgrability of ∇*b*, we must have *C*=0, and so *b* is in fact Lipschitz-continuous at the origin, which justifies Eq. 20. Lastly, boundedness of *F*[*b*], Eq. 45, Lipschitz continuity of *b* and decay of *b*_RBA_−1 at infinity yield *b*(*r*)→1 as *r*→∞.

